# Biosurfer for systematic tracking of regulatory mechanisms leading to protein isoform diversity

**DOI:** 10.1101/2024.03.15.585320

**Authors:** Mayank Murali, Jamie Saquing, Senbao Lu, Ziyang Gao, Ben Jordan, Zachary Peters Wakefield, Ana Fiszbein, David R. Cooper, Peter J. Castaldi, Dmitry Korkin, Gloria Sheynkman

## Abstract

Long-read RNA sequencing has shed light on transcriptomic complexity, but questions remain about the functionality of downstream protein products. We introduce Biosurfer, a computational approach for comparing protein isoforms, while systematically tracking the transcriptional, splicing, and translational variations that underlie differences in the sequences of the protein products. Using Biosurfer, we analyzed the differences in 32,799 pairs of GENCODE annotated protein isoforms, finding a majority (70%) of variable N-termini are due to the alternative transcription start sites, while only 9% arise from 5’ UTR alternative splicing. Biosurfer’s detailed tracking of nucleotide-to-residue relationships helped reveal an uncommonly tracked source of single amino acid residue changes arising from the codon splits at junctions. For 17% of internal sequence changes, such split codon patterns lead to single residue differences, termed “ragged codons”. Of variable C-termini, 72% involve splice- or intron retention-induced reading frameshifts. We found an unusual pattern of reading frame changes, in which the first frameshift is closely followed by a distinct second frameshift that restores the original frame, which we term a “snapback” frameshift. We analyzed long read RNA-seq-predicted proteome of a human cell line and found similar trends as compared to our GENCODE analysis, with the exception of a higher proportion of isoforms predicted to undergo nonsense-mediated decay. Biosurfer’s comprehensive characterization of long-read RNA-seq datasets should accelerate insights of the functional role of protein isoforms, providing mechanistic explanation of the origins of the proteomic diversity driven by the alternative splicing. Biosurfer is available as a Python package at https://github.com/sheynkman-lab/biosurfer.

## INTRODUCTION

Through the isoform diversifying mechanisms of alternative transcription, splicing, and polyadenylation, nearly every human gene can produce multiple protein products, with ∼20K genes giving rise to at least 180K annotated isoforms (Frankish et al. 2023). The pathway from gene to protein is marked by several regulatory mechanisms that are highly tuned across development and cell states, with disruption of this regulation producing aberrant isoforms that lead to pathophysiological states such as cancer and cardiovascular disease (Cooper et al. 2009). Hence, approaches are needed to systematically characterize the upstream regulatory causes and functional impacts of such protein isoform sequence changes.

Transcript and, by extension, protein isoform diversity may now be globally characterized at great depth for individual samples (Glinos et al. 2022; Reese et al. 2023). Transcript diversity can be readily characterized by long read RNA-Seq, which employs single molecule sequencing of individual cDNA or RNA molecules to determine the sequence across the entire length of spliced transcripts (Sharon et al. 2013; Workman et al. 2019; PardolPalacios et al. 2021; Tian et al. 2021; Joglekar et al. 2023), with platforms from Oxford Nanopore and PacBio being most commonly used (Clarke et al. 2009; Eid et al. 2009). Long read RNA sequencing captures long range connectivity between multiple exons of a transcript and can reveal complex splice patterns unattainable by short read sequencing (De PaolilIseppi et al. 2021), including dependencies across distal splicing events (Anvar et al. 2018) and alternative 5’ and 3’ transcript usage. Given this readily characterized complexity afforded by long read sequencing, a natural question is the extent to which such variations of the transcriptome lead to functional effects of the proteome. Towards this goal, a necessary step is defining the potential proteome. Both our group and others have reported methods in which long-read-derived transcript sequences serve as templates for predicting full-length protein isoform sequences, thereby providing a global snapshot of potential protein isoforms expressed in a particular biological condition (Miller et al. 2022; Veiga et al. 2022; Abood et al. 2023).

The complexity of alternative splicing (AS)—which for practical purposes in this manuscript we define here as all transcriptional variations, including alternative transcription start sites (TSS) and transcription termination sites (TTS)—can be observed and characterized at different levels: RNA transcript, open reading frames (ORFs), and finally protein sequences (ReixachslSolé and Eyras 2022). The complex interplay between the changes occurring to AS variants and their ORF and protein products are hard to characterize and quantify. Changes in mRNA sequence may lead to non-linear or traditionally untracked variations. For example, a subtle splicing event could lead to a reading frame shift and thus to more dramatic changes to the C-terminus of the protein than the originating small change at the RNA level would suggest. Or, AS could occur at codon boundaries, leading to altered amino acid (AA) identities of codons that technically overlap in genome-space but are differentially “split” across exon-exon junctions. And complex interplay may also be observed between transcriptional variations and ORF choice, as alternative 5’ transcription or AS could lead to differentially availability of initiator codons, delimiting start codon choice co-translationally.

We argue that rather than being an esoteric exercise, the ability to characterize all potential interplay of RNA-protein variation is critical for fully elucidating the transcript and proteomic diversity encoded within long-read RNA-seq datasets. As long-read RNA-seq approaches are increasingly adopted in large-scale studies of hundreds of samples (Glinos et al. 2022; Reese et al. 2023) and are maturing into stable tools being adopted by the community (PardolPalacios et al. 2021), extracting all sources of biological molecular diversity is critical. Such interplay cannot be characterized by comparison of isoforms using just one modality, such as transcript-focused annotation tools like SQANTI or Matt (Tardaguila et al. 2018; Gohr and Irimia 2019), or conventional protein sequence alignment tools like ClustalW (Chenna et al. 2003) or BLAST (Altschul et al. 1990). Recently, multi-modal comparisons have been reported. For example, ORFanage is an approach for large-scale annotation of ORFs across predicted transcripts in the CHESS database the main focus being optimal selection of ORFs based on protein-alignments (Varabyou et al. 2023). Other tools geared towards the phylogenetics community have developed frame-aware alignment in which the alignment scoring system is includes penalties for frame-shift-inducing gaps <u>(Evans and Loose 2015; Ranwez et al. 2011; Jammali et al. 2022)</u>. However, these tools do not comprehensively elucidate the interplay between transcript and protein variation.

To characterize how AS impacts protein sequence, several bioinformatic tools and databases have been developed, such as VastDB, ASPicDB, ExonOntology, and DIGGER (Martelli et al. 2011; Tapial et al. 2017; Tranchevent et al. 2017; Louadi et al. 2021). Tools such as tappAS and IsoTV annotate how protein isoform sequence and potential functional differences (de la Fuente et al. 2020; Annaldasula et al. 2021). For example, tappAS is a Java application for quantifying differential isoform usage but also to functionally annotate such isoforms, using the module IsoAnnot (de la Fuente et al. 2020). IsoAnnot maps protein features (e.g., Pfam domains) across isoforms of a gene, and determines how splicing leads to partial or full removal of protein features, indicating potential changes to molecular functions. Despite existence of these tools, of need is the ability to systematically capture all possible effects to protein isoforms, with the accompanying information of the underlying complex RNA-protein relationships.

We developed Biosurfer, a computational pipeline that tracks simultaneously the changes at all three levels, to understand the impact of alternative splicing on transcriptome, ORFeome, and proteome diversity. Biosurfer computes details not immediately apparent from genome annotation files or manual inspection in genome browsers, such as how between isoforms of the same gene the frame of translation and codon topology influences amino acid sequence identity changes. In order to accomplish this multi-layered comparison, a genome is used as a “scaffold” to exactly position all nucleotides, codons, AA elements, and associate with each element local and context-dependent attributes. The resulting data structure of three distinct yet interlinked layers inform on the upstream biological mechanisms leading to protein sequence changes. To demonstrate the utility of Biosurfer, we globally characterize variation across protein isoforms in the reference human annotation (GENCODE) and protein isoforms predicted from long-read RNA-seq of a human stem cell. This characterization includes comprehensive elaboration of all sources of N-terminal, internal, and C-terminal protein variations observed, highlighting mechanisms of RNA-protein interplay.

## METHODS

### Biosurfer package for isoform analysis and visualization

Biosurfer is a computational pipeline that performs a multi-layered comparison between a pair of isoforms, in which differences at three different levels: RNA (nucleotides, nt, ORF (codon), and protein (AA residues, AA) are simultaneously tracked. The developed data structure enables not only the comparison of AAs, but tracking frameshifts, patterns of codon splitting at junctions, and the attendant upstream nucleotide and codon differences that explain AA changes. Such tracking aids in the systematic annotation of explanatory mechanism(s) underlying AA residue changes, such as whether a substitution of a stretch of AA residues is due to alternative splicing or a frameshift (**Supplementary Figure S1**).

The Biosurfer pipeline is organized into the following three stages (**Figure 1)**. First, an SQLite database is populated with detailed information on each isoform at the transcript, ORF, and protein product levels. For each isoform, the required inputs are (i) a transcript FASTA, (ii) a protein FASTA, and (iii) a matching GTF with both Exon and CDS features. The inputs can either be extracted from the reference annotations (*e.g.*, GENCODE (Harrow et al. 2012)), or they can be user-defined (*e.g.,* predicted protein isoform sequences from long-read RNA-seq data (Miller et al. 2022)by using ORF callers such as CPAT (Wang et al. 2013) or Transdecoder (Haas et al. 2013)). Second, multilayered isoform-level alignments are generated, and each alignment is represented as three key data structures: t-blocks, c-blocks, and p-blocks corresponding to the view of the aligned isoforms at the transcript, codon sequence, and protein sequence levels, respectively (**Supplementary Figure S2**). Third, all information is summarized in tabular format, to facilitate analysis of the transcription-level and codon-level mechanisms driving the proteomic diversity. In addition, a visual representation of such mechanisms integrated with the protein-level view of the isoforms can be output as png files.

**Figure 1:**
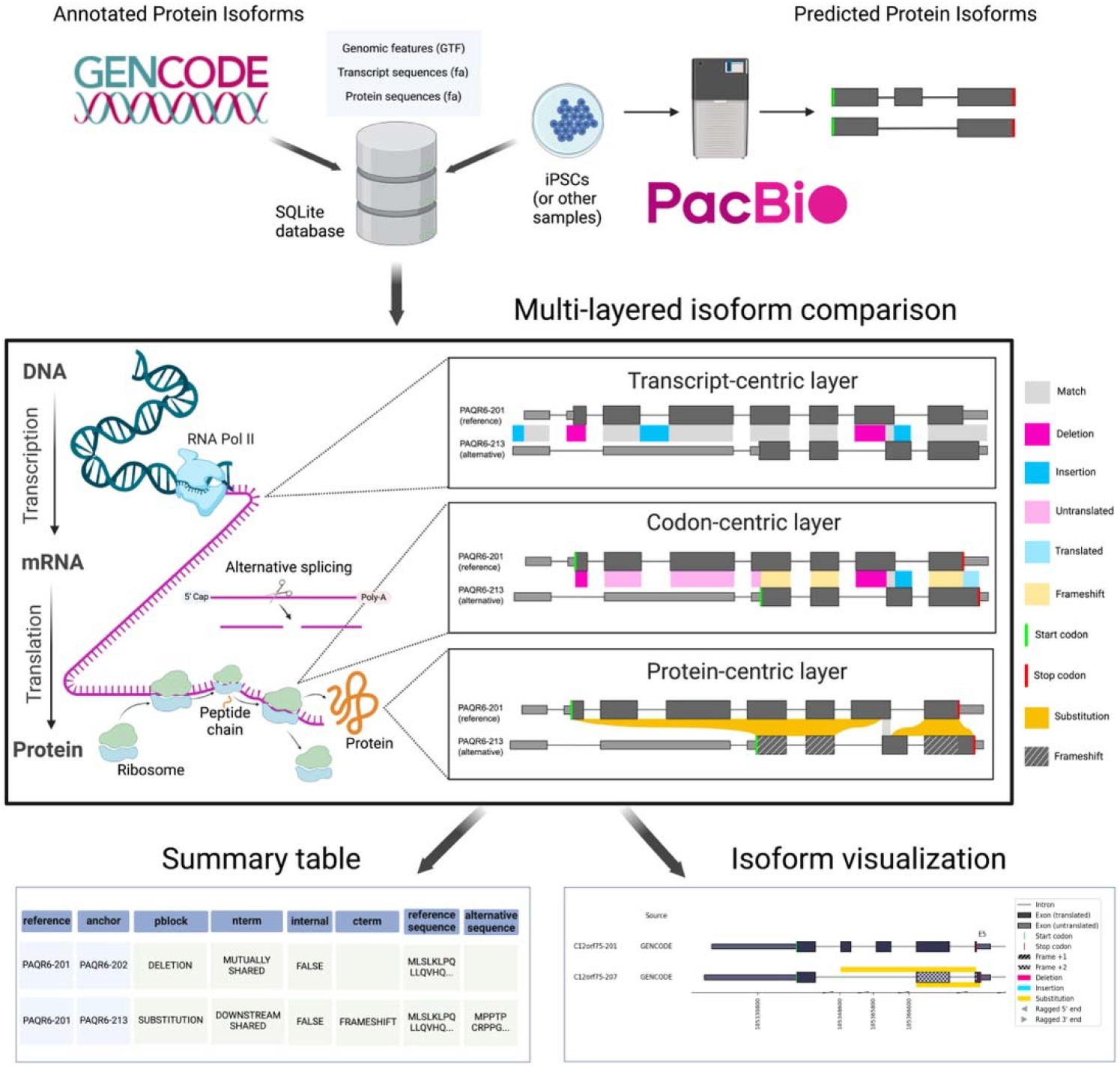
Biosurfer for analysis and visualization of protein isoform sequence differences. Biosurfer analyzes protein isoforms from reference annotations (e.g., GENCODE) or proteins predicted from long-read RNA-seq data. Analysis initiates with the creation of an SQLite database is populated with isoform-relevant elements. Biosurfer performs a multi-layered comparison of transcript-, codon-, and protein-level differences between pairs of protein isoforms. Variable regions as well as their associated annotations are output in tabular format and visualization files, which includes protein-relevant details such as the frame of translation. Note that the terms “Match”, “Deletion”, etc. represent very different comparisons depending on the biological layer. GTF=Gene Transfer Format; iPSCs=induced pluripotent stem cells.

### Data structures for presenting transcript-, ORF-, and protein-level isoform alignments in Biosurfer

Biosurfer compares transcript-, ORF-, and protein-level sequences for each gene. One isoform is selected as reference and the other isoforms are denoted as alternative. When aligning each alternative isoform to the reference, the matched regions are referred to as matched **blocks**. The remaining regions on each sequence that are not matched blocks are referred to as unmatched blocks. See **Supplementary File 1** for step-by-step schema of the comparison process, which is summarized below.

#### Transcript blocks (t-blocks)

T-blocks represent subsegments of the transcript sequence that are shared or unique to the reference or alternative isoforms. Transcript-level differences are determined by analyzing the transcript-to-genome coordinates alignment (GTF file) provided as an input by the user. Specifically, Biosurfer defines the aligned exonic regions that are shared or unique to each isoform. The resulting ranges, called *t-blocks*, are categorized as Match, Deletion, or Insertion t-blocks. Deletion or Insertion t-blocks are further annotated with the associated biological mechanism leading to the transcript nucleotide change, *e.g.*, alternative transcriptional start site, splicing event, or polyadenylation. The alternative splicing events are then further classified into four basic types: retained intron, alternative donor, alternative exon, or alternative acceptor.

#### Codon blocks (c-blocks)

C-blocks represent a codon-centric layer defined through the comparison of the protein coding regions of transcripts, *i.e*., open reading frames (ORF), between two isoforms. The c-block data structure is the most complex, but critical, layer in Biosurfer that connects information between the transcript and protein layers.

For ultimate granularity and precision, ORFs are compared based on the alignment of codons that overlap in the genome space, in which one codon in the reference isoform is compared with another codon in the alternative isoform (see **Supplemental Methods** for additional details). First, codons across the two isoforms are “**paired**” based on their mutual positions and base overlap: in a basic scenario, the two codons are identical and their positions match (**Table1**, **Supplemental Figure S3A**); in a more complex scenario, the codons are split and only partially overlapped (1 or 2 bases, see **Table1**, **Supplementary Fig S3B** and **C**). In all other cases, the codons will be designated as “unpaired” (**Table1**,

**Supplementary Figure S3B**). To keep track of unpaired codons within the data structure, they are linked to a “placeholder” codon, which serves to maintain consistency of the c-block structures. Once aligned, each single codon pair is categorized with respect to multiple attributes that explain their relationship.

Overall, the paired and unpaired codons are classified into 9 categories based on their translation status, frameshift status, and codon topology (**Table 1**)

**Table 1:**
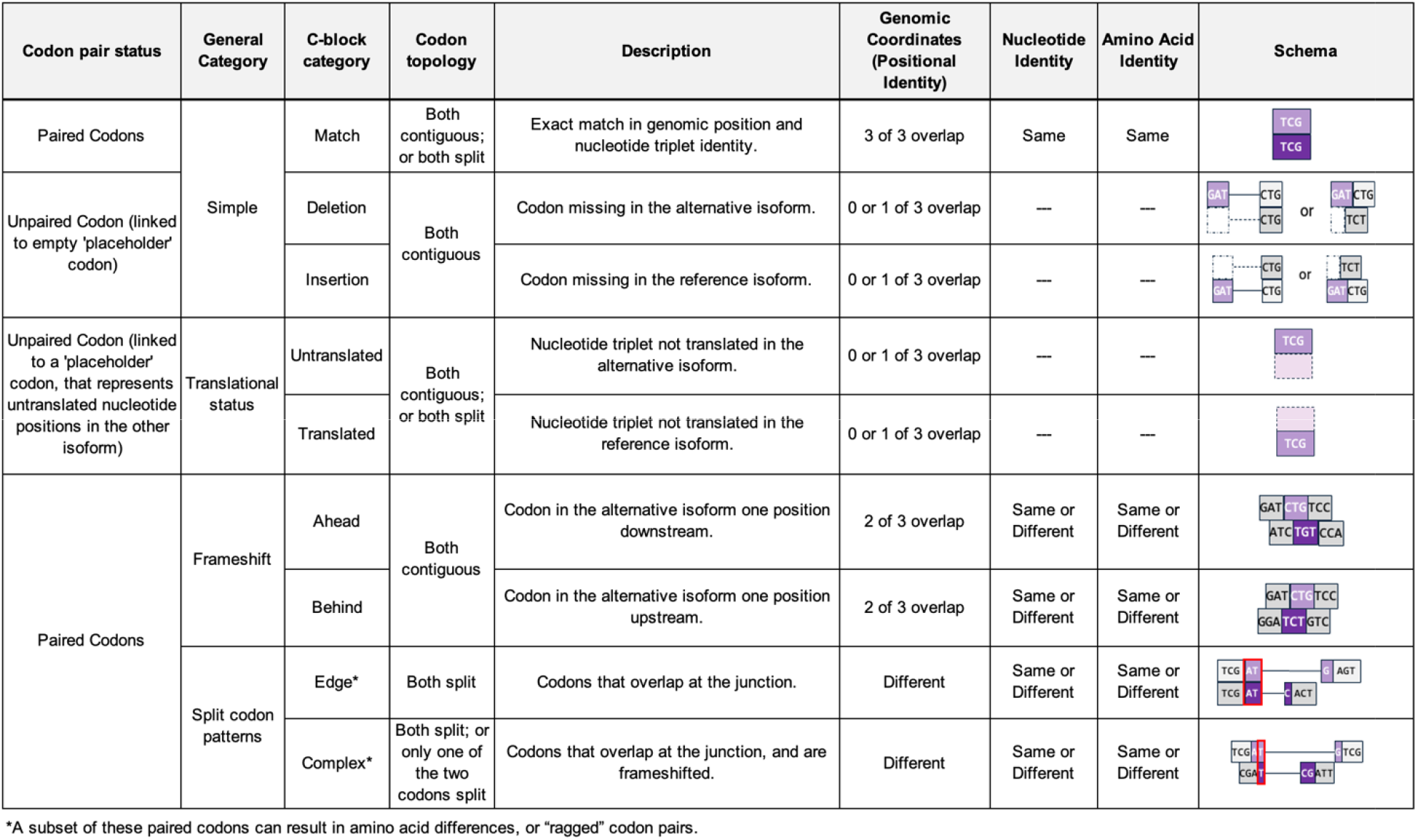
Categories of overlapping codon pairs that are the basis for codon blocks (c-blocks).

| Codon pair status | General Category | C-block category | Codon topology | Description | Genomic Coordinates (Positional Identity) | Nucleotide Identity | Amino Acid Identity | Schema |
| --- | --- | --- | --- | --- | --- | --- | --- | --- |
| Paired Codons | Simple | Match | Both contiguous;<br>or both split | Exact match in genomic position and nucleotide triplet identity. | 3 of 3 overlap | Same | Same |  |
| Unpaired Codon (linked to empty 'placeholder' codon) |  | Deletion | Both contiguous | Codon missing in the alternative isoform. | 0 or 1 of 3 overlap | --- | --- |  |
|  |  | Insertion |  | Codon missing in the reference isoform. | 0 or 1 of 3 overlap | --- | --- |  |
| Unpaired Codon (linked to a 'placeholder' codon, that represents untranslated nucleotide positions in the other isoform) | Translational status | Untranslated | Both contiguous;<br>or both split | Nucleotide triplet not translated in the alternative isoform. | 0 or 1 of 3 overlap | --- | --- |  |
|  |  | Translated |  | Nucleotide triplet not translated in the reference isoform. | 0 or 1 of 3 overlap | --- | --- |  |
| Paired Codons | Frameshift | Ahead | Both contiguous | Codon in the alternative isoform one position downstream. | 2 of 3 overlap | Same or Different | Same or Different |  |
|  |  | Behind |  | Codon in the alternative isoform one position upstream. | 2 of 3 overlap | Same or Different | Same or Different |  |
|  | Split codon patterns | Edge* | Both split | Codons that overlap at the junction. | Different | Same or Different | Same or Different |  |
|  |  | Complex* | Both split; or only one of the two codons split | Codons that overlap at the junction, and are frameshifted. | Different | Same or Different | Same or Different |  |

\*A subset of these paired codons can result in amino acid differences, or “ragged” codon pairs.

#### Protein blocks (p-blocks)

P-blocks represent a protein-centric layer defined through t- and c-block guided comparisons of two protein isoforms, thus focusing solely on the relationships between the AA residues of the respective protein isoforms. This decision was motivated by the fact that in most cases, functional differences exhibited by an alternative protein isoform are attributable to differences in protein rather than nucleotide sequences between the alternative and reference isoforms, and these differences could be conceptually decoupled from the knowledge of the upstream (ORF- or transcript-level) residue-altering mechanisms.

The *p-blocks* for a pair of isoforms are determined by translating c-blocks defined at the previous level. The resulting data structure represents subsegments of contiguous stretches of AA residues from each protein sequence. Each subsegment could be as short as a small translated portion of exon (even a single residue, in the case of NAGNAG splicing), but it can also span multiple exons of a gene. Because the actual matching happened at the two previous levels (t-blocks and c-blocks), no alignment is required at the p-block level. The resulting protein subsegments can be either (1) fully matched (100% sequence identity, across all residues in the subsegment), or (2) mismatched, where a subsegment in one protein sequence will be aligned against a gapped region in the other sequence. Thus, a p-block represents either a matched pair of protein subsequences or a subsequence matched against a gapped region. P-blocks are then classified as Match, Insertion, Deletion, or Substitution changes. Substitution p-blocks must arise from a combination of insertion/deletion/frameshift events found at the c-block level.

Overall, these p-block changes are agnostic to the upstream mechanisms, *e.g*., at the transcript or ORF levels; however, at the same time, the corresponding upstream mechanisms can be retrieved and analyzed within Biosurfer, unlike using traditional protein aligners.

### Analysis of GENCODE isoforms using Biosurfer

The principal and alternative isoforms were defined using APPRIS annotation of genes in GENCODE (Rodriguez et al. 2013). First, the set of APPRIS isoforms is identified for each gene from the input genome data (GENCODE v42, basic annotation (Frankish et al. 2021)) by extracting the key transcript features, such as ‘transcript_id’, ‘transcript_name’, and the associated APPRIS ‘tag’ (Rodriguez et al. 2013). Second, the transcript’s APPRIS status is determined as ‘principal’, ‘alternative’, or ‘none’, based on first rank-ordering of transcripts based on APPRIS tag and setting the transcript with the highest APPRIS value as ‘principle’ and all other transcripts as ‘alternative’. We did not consider further genes with only one annotated isoform and lacking an APPRIS tag (‘none’). Subsequently, we compiled a structured dataset for each transcript, encompassing identifiers, gene associations, strand orientation, and APPRIS status. During the annotation, the strand orientation is accounted for when necessary.

### Long read RNA-seq analysis of WTC-11 cell line

Total RNA from WTC-11 cells was extracted using the RNeasy Kit (Qiagen) and analyzed on an Agilent Bioanalyzer. We observed a RNA concentration of 30 ng/uL with the RNA Integrity Score (RIN) of 9.9. As described previously, (Mehlferber et al. 2022) cDNA was synthesized from the extracted RNA and the Iso-Seq Express Kit SMRT Bell Express Template prep kit 2.0 (Pacific Biosciences) was used on a Sequel II system to obtain long-read sequence information and output Circular Consensus (CCS) reads. Data is available at the Sequence Read Archive: SRR18130587 and previously published (de Souza et al. 2022).

We analyzed the WTC-11 data with a proteogenomics Nextflow pipeline we previously developed (Miller et al. 2022) (Mehlferber et al. 2022). The output CCS reads from long-read sequencing were processed with Iso-Seq3 and SQANTI3 (version 1.3) for transcript isoform classification and quality assessment. The CPAT(Wang et al. 2013) algorithm was used to predict Open Reading Frames (ORFs), which were grouped into protein isoforms.

Biosurfer is implemented as a Python package freely available at GitHub repository: https://github.com/sheynkman-lab/biosurfer. The analysis code is available at https://github.com/sheynkman-lab/biosurfer_analysis. All necessary input files and intermediate and final output files from Biosurfer analysis are uploaded to Zenodo at: https://zenodo.org/records/10822882.

## RESULTS

### Characterization of altered protein regions in the human annotation (GENCODE)

Here, we demonstrate the utility of Biosurfer through a genome-wide analysis of protein isoforms in the GENCODE annotation (basic annotation, version 42 (Frankish et al. 2021)). We analyzed 35,083 reference-alternative protein isoform pairs across 11,815 genes (**Figure 2A**). Each pair consists of the reference protein isoform for a gene—the APPRIS principal isoform (Rodriguez et al. 2013)—and an alternative protein isoform. The number of isoform pairs correspond to the number of “alternative” isoforms for a gene (**Supplementary Table S1**). Genes lacking an APPRIS annotation (8,166 of 19,981) were excluded from the analysis (**Supplementary Table S2**).

**Figure 2:**
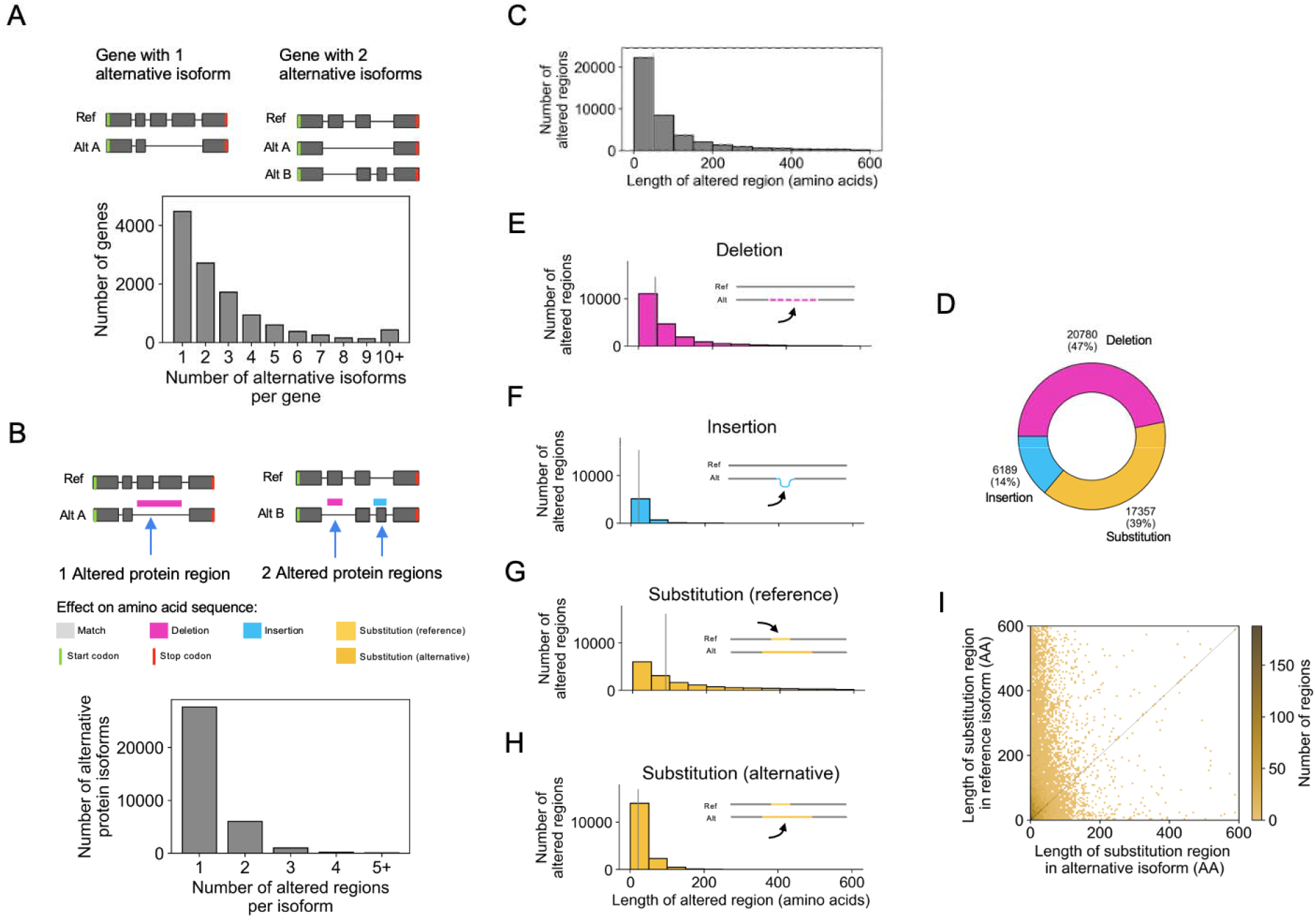
Characterization of altered protein regions (Biosurfer p-blocks) across the GENCODE annotated human proteome. (A) Schematic of genes with one or two alternative protein isoforms and distribution of the number of alternative protein isoforms per gene. (B) Schematic of the altered protein regions (here, highlighted in pink and blue), displayed relative to the underlying transcript structures. Th proteome-wide distribution of the number of affected protein regions observed per alternative isoform. (C) Distribution of the length of altered protein regions across the annotated proteome. Differences greater than 600 AAs are not included (2,347 cases, 5.3% of the data). These altered protein region include cases of 1) deleted regions, 2) inserted regions, and 3) the region (in the reference isoform) in which one polypeptide subsegment is substituted for another. In other words, this distribution (C) represents an aggregation of the distributions shown in panels E-G. (D) Fraction of altered protein regions affected by insertions, deletions, and substitutions. (E-H) Distribution of the lengths of altered protein regions for (E) deletions, (F) insertions, (G) substituted region in the reference isoform, and (H) substituted region in the alternative isoform. (I) Comparison of the lengths of altered regions in the reference versus alternative isoforms for substitutions.

Globally, we found a total of 44,326 altered protein regions with an average of 1.3 altered regions per isoform. Note that we are using the term altered protein region interchangeably with p-block (see **Methods**). Altered protein regions are contiguous regions of altered protein sequence relative to the “reference” (i.e., APPRIS principal) protein, which includes p-block Insertions, Deletions, and Substitutions. A majority (79%, 27,672 of 35,083) of isoform pairs contain a single altered region (**Figure 2B**). Still, 7,411 alternative isoforms contain two or more discontinuous altered regions. Notably, some isoform pairs exhibited an extreme number of regions. For example, 14 regions are found for proteins of *DNAH14* (Reference: DNAH14-220, Alternative: DNAH14-211), which is explained by its extremely large number of exons (86 exons for DNAH14-220).

Among the 44,326 altered protein regions (**Figure 2C**), the median number of affected amino acids (lost or gained due to a protein insertion, deletion, or substitution) is 49 AA, with the first and third quartiles containing 21 AA and 128 AA, respectively. Among the altered regions, 14% (6,189) are insertions, 47% (20,780) are deletions, and 39% (17,357) are substitutions (**Figure 2D**). Full annotations for these altered regions at the protein and codon-level, representing the Biosurfer output for protein-blocks (p-blocks) and codon-blocks (c-blocks) can be found in **Supplementary Table S3**, **S4**.

The lengths of p-block insertions tend to be shorter than the length of deletions (p < 2.2e-16, Mann-Whitney U test) or substitutions (p < 2.2e-16, Mann-Whitney U test) (**Figure 2E-H**). Since the deletion or insertion status of a protein region is dependent on which isoform is denoted as the reference, this trend may reflect the tendency that longer isoforms are more likely to be defined as the “reference” isoform, which may be biologically driven or influenced by genome annotation guidelines (Rodriguez et al. 2013; O’Leary et al. 2016; The UniProt Consortium 2017; Frankish et al. 2021; Varabyou et al. 2023). We found a similar trend for p-block substitutions; the lengths for substituted regions tended to be longer for the sequence affected in the reference (**Figure 2G**) versus the alternative (**Figure 2H**) isoform; although, for 3,263 (18% of cases), the affected regions in alternative isoforms can be longer (**Figure 2I**).

### Analysis of N-terminal protein variations

Variations in the N-termini are found in 28% (12,504 of 44,326) of all possible reference-alternative isoform pairs, corresponding to 5,872 genes (**Supplementary Table S5**). We examined for the N-terminal variations the explanatory mechanism, including alternative transcription start sites (TSSs), alternative splicing, and alternative translation initiation sites (TISs).

All cases of variable N-termini involve two initiation codons (AUGs), one upstream and one downstream, relative to the genome. A major category we first observed are those in which the N-terminus is different due to start codons that are mutually exclusively present across the two transcript isoforms (**Figure 3A**). Specifically, the start codon present in the reference isoform is absent from the alternative isoform, and vice versa. These “mutually exclusive start codons” or MXS were observed for 3,123 reference-alternative isoform pairs (**Figure 3A**). MXS may arise either from an alternative TSS or from alternative splicing in the 5’ UTR. Strikingly, we found that nearly all (99%, 3,097 of 3,123) cases are caused by alternative TSS usage (**Figure 3A**, hatched region of the bar), with only a small, but non-zero, fraction (1%, 26 of 3,123) of MXS cases arise from splicing of the 5’ UTR, in which splicing regulation is influencing N-terminal usage. An example of TSS-driven MXS for *PRKACA* is shown in **Figure 3B** (pair, PRKACA-201 and PRKACA-202).

**Figure 3:**
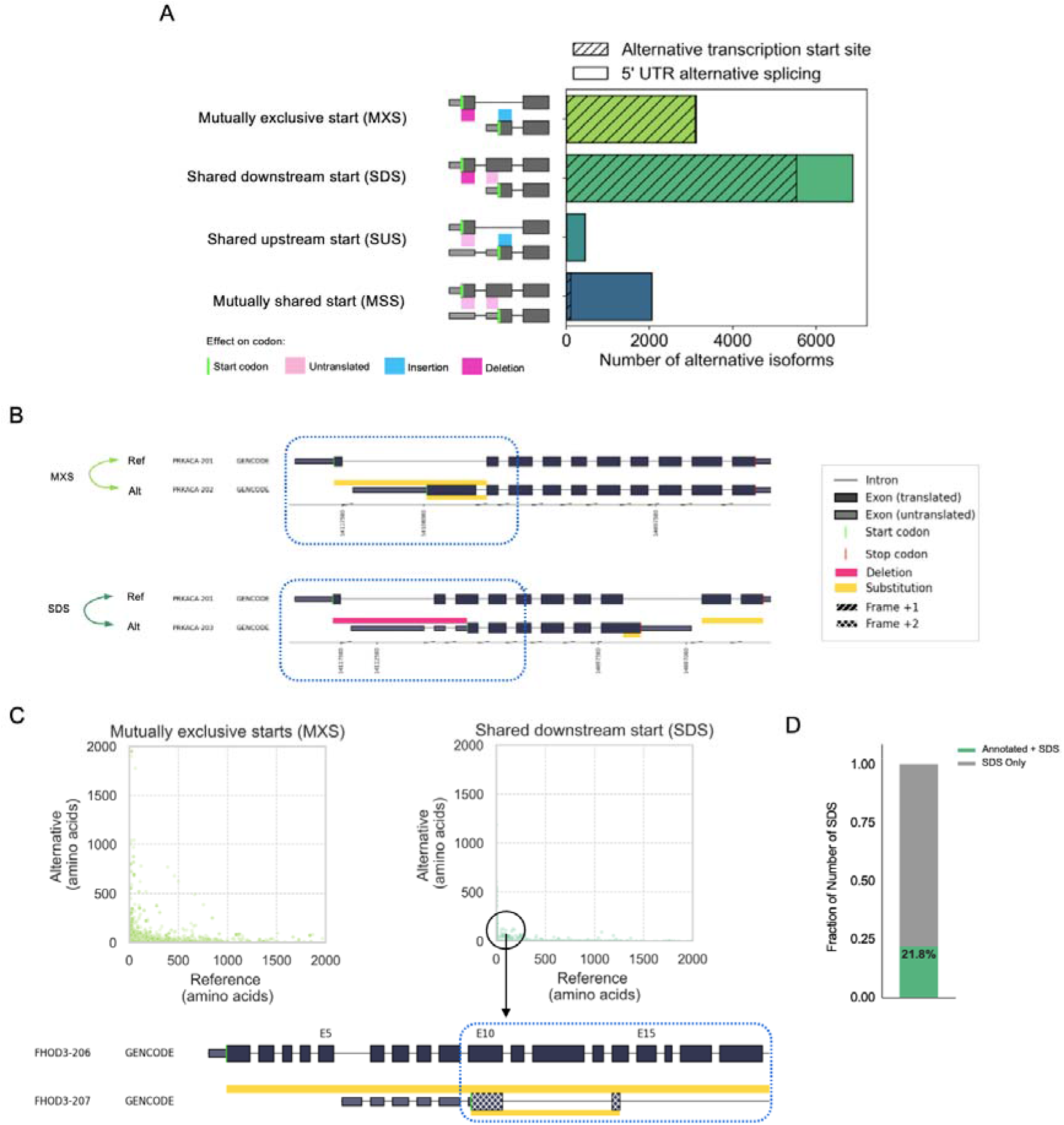
Analysis of mechanisms underlying variable N-terminal proteins across the GENCODE annotated human proteome. (A) Distribution of the types of alternative N-terminal regions, classified based on presence and translational status of the start codon. Hatches denote the fraction of alternative N-terminal regions associated with alternative transcription start sites, as opposed to 5’ UTR alternative splicing. (B) Biosurfer output of altered N-terminal regions for *PRKACA* gene that undergo MXS (light green arrows) and SDS (dark green arrows). In the example of MXS, the yellow bars above and below PRKACA-202 (alternate) transcript indicate the N-terminal ranges that differ between the reference and alternative isoform. The yellow bar above PRKACA-202 shows the N-terminal protein sequence that is specific to the reference (PRKACA-201) and the bar below the transcript indicates the N-terminal region specific to the alternative isoform. The Biosurfer bars span the intronic lines between exons, but intronic regions do not contribute to the protein sequence differences. In the example of SDS, the pink Biosurfer bar above the transcript of PRKACA-203 represents the range of transcript sequence that is translated in the reference, but not translated in the alternative isoform. (C) Scatterplot of the length of affected N-terminal variation in the reference versus alternative, faceted by mutually exclusive starts (MXS) or shared downstream start (SDS) status. An interesting case of an SDS leading to unique N-terminal sequence in the reference is caused by usage of a different frame at the initiation of translation, with an example shown for isoforms of the gene *FHOD3*. (D) Fraction of shared downstream starts (SDS) caused by hybrid exon swaps.

The second category is when the upstream start codon is transcribed in only one of the two isoforms, but the downstream start codon is present in both transcripts. We refer to this scenario as shared downstream starts (SDS), of which there were 6,878 cases (**Figure 3A**). Like with cases of MXS, SDS arises primarily from alternative TSS usage (80% of cases) versus 5’ UTR alternative splicing (20% of cases).

MXS and SDS are common patterns underlying alterations of the N-termini or protein, driven by differential availability of initiator codons in the mature transcript. We asked if there may be differences in length of such N-terminal alterations for SDS versus MXS events. Measuring the differential length of the affected N-terminal regions between reference-alternative isoform pairs, we found that, on average, SDS tends to affect a greater proportion of protein length, as compared to MXS (**Figure 3C**, **Supplementary Figure S4**; p = 2.8e-148, Mann-Whitney U test). The larger differences in length driven by SDS could be explained by cases in which transcription is initiated from internal sites of the gene, giving rise to an ORF that corresponds to a subsequence of the ORF in the other isoform, theoretically producing a truncated C-terminal-containing subsequence of the full-length protein.

Recently, a mechanism related to SDS was described in which internal exons (not the 5’ most exon, first exon, of a transcript) in one transcript can be immediately downstream of a DNA element of novel promoter activity and thus serve as the first transcribed exon in other isoforms (Fiszbein et al. 2022). Such so-called hybrid exons thus can operate as both sites of transcription initiation and alternative splicing, in effect, swapping their roles depending on the regulatory context. Of the cases of SDS, we observed that ∼22% correspond to these hybrid exon swaps (**Figure 3D**, **Supplementary Table S6**). The functional consequences of hybrid exon usage are not well understood; however, one potential function could be the production of protein with a truncated N-terminus, which could remove signal peptide sequences or binding domains (Kelemen et al. 2013).

In addition to MXS and SDS, wherein start codon availability is controlled through differential transcription, we also observed many cases in which both upstream and downstream start codons co-occur in one or both transcript isoforms of a pair. In these cases, the choice of start codon may be influenced by co-translational regulation, e.g., ribosome initiates translation at alternative initiation sites (altTIS).

For 2,054 cases, we found altTIS, which we classified as instances of a mutually shared start (MSS) codon. We also found 449 cases in which the upstream start codon is present in both isoforms, but the downstream start codon is only present in one isoform.

In such cases of shared upstream start (SUS) (**Figure 3A**), the explanatory co-translational mechanism is not as clear, based on the ribosomal scanning model of translation initiation (Kozak 1978). The upstream start codon present in both isoforms would need to be bypassed by the ribosome only in the alternative isoform. Therefore, some of the SUS annotations may need to be validated or may be erroneous ORF calls, as early ORF prediction workflows attribute higher scores to longer ORFs (Wang et al. 2013; Varabyou et al. 2023), under-annotating ORFs that utilize the upstream (annotated) start codon but is much shorter than the reference due to a reading frame shift (Wang et al. 2013).

### Analysis of internal protein variations

The internal regions of the protein isoforms account for 43% (19,263 of 44,326) of possible reference-alternative isoform pairs, corresponding to 6,673 genes (**Supplementary Table S7**). A large majority of these regions (80%, 15,444 of 19,263) are caused by single simple splicing events: exon skipping, alternative acceptor, alternative donor, or an in-frame retained intron. As expected, exon skipping events are most numerous making up 9,505 of cases. In terms of the general effect on protein sequence, most altered regions (70%, 13,453 of 19,263) lead to a deletion or removal of AA residues (**Figure 4A**), and, again, in such cases, exon skipping is most common (51%, 6,908 of 13,453).

**Figure 4:**
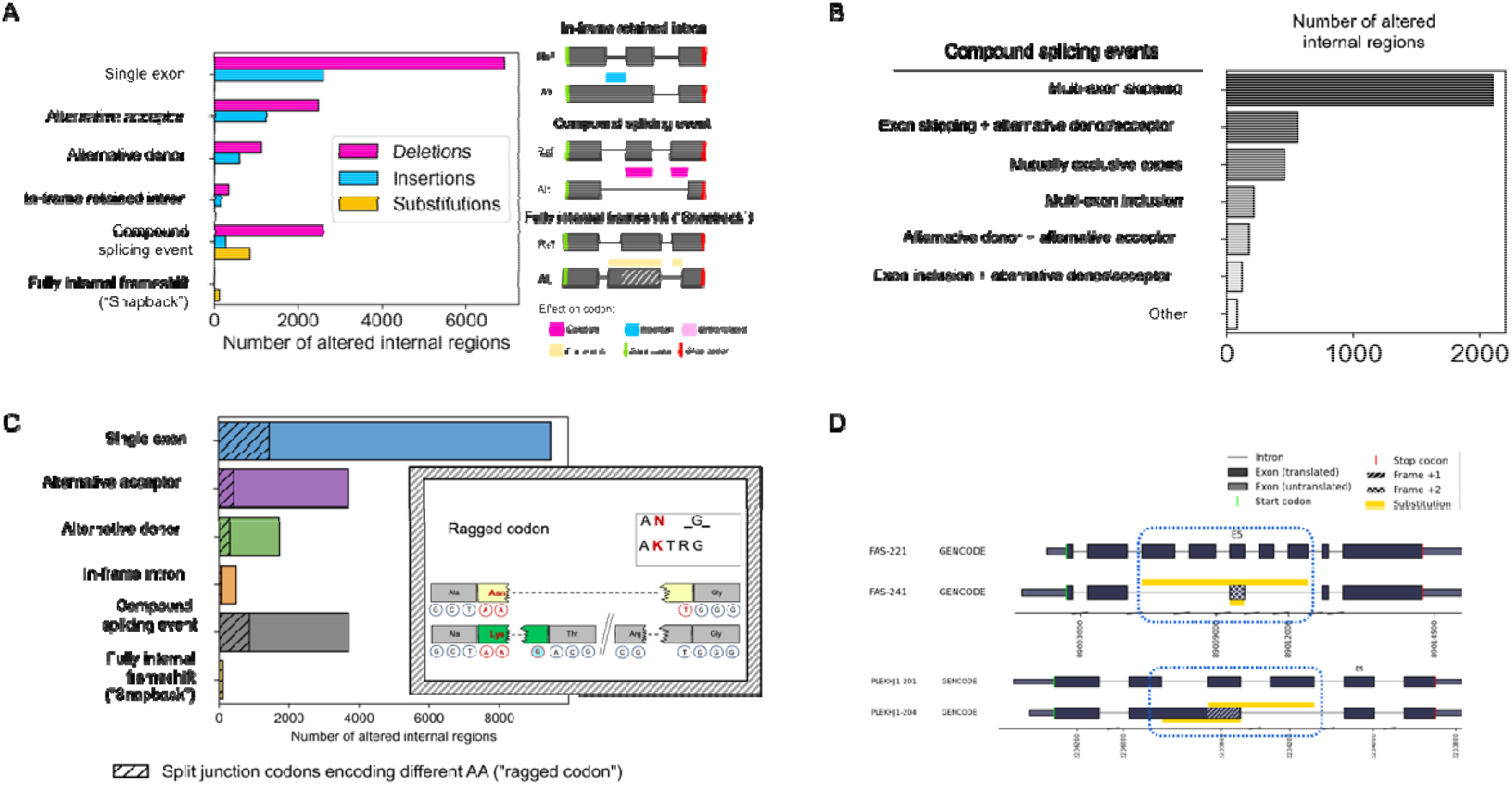
Analysis of internal protein altered regions across proteins in GENCODE. (A) Frequencies of the categories of the splicing mechanism underlying internal protein sequence changes, split by their protein-coding impact (Deletion, Insertion, Substitution). All sequence regions involve a reference-alternative isoform pair. (B) Frequency of compound splicing events across the altered internal regions. (C) Proportion of each internal protein region type for which there exists a split codon pattern near its boundaries that would cause a single amino acid difference, or “ragged” codon. (D) Examples of successive frameshifting (snapback” frameshift) that leads to an affected protein region that is wholly internal to the protein, for genes *FAS* and *PLEKHJ1*.

Going beyond simple splicing events, we observed that 19% of variable internal regions (3,701 of 19,263) were associated with multiple events, which we refer to as compound, or linked, splicing events. We found that 56% (2,099 of 3,701) compound events involve multi-exon skipping, the rest being combinations of alternative donor/acceptor sites with exon skipping/inclusion (**Figure 4B**).

### Complex nucleotide to amino acid relationships that affect internal protein sequence

Previous studies of the impact splicing on proteins have typically focused on cases in which differential splicing of transcript regions directly corresponds to changes in protein sequence (ReixachslSolé and Eyras 2022). However, in many instances, there is not a simple one-to-one relationship between nucleotides in a transcript and the corresponding amino acid identities in the encoded protein. Such complexity alters the protein sequence in non-intuitive ways. Using the detailed codon tracking afforded by Biosurfer, we systematically characterized the protein-level impact of variations not commonly described: codons that span junctions and unusual reading frame shifts.

To characterize differentially split codons, we examined all paired codons (see c-block section in **Methods**) that are split across junctions and determined the identity of the associated AAs (**Supplemental Figure S3**). Across all internal altered protein regions, we found 17% (3,213 of 19,263) of regions that are flanked by one or more split codon pairs that encode different amino acid residues (**Figure 4C**, also see **Table 1**). These so-called ragged codons affect a single residue and are always adjacent to an altered protein region. Interestingly, while split codons are not particularly enriched by splice event type (Chi-square test: p-value = 5.54e-99), we found that ragged codons are more frequently found in protein insertions compared to deletions or substitutions (Chi-square test: p-value = 4.11e-106).

For a majority of splice-driven frame shifts, the shifted frame is maintained to the end of the protein, leading to a protein variation affecting the C-terminus. Such frameshifted proteins could lead to truncated proteins or destabilization of the transcript via mechanisms such as nonsense mediated decay (NMD). Surprisingly, we found an uncommonly characterized pattern of successive reading frame shifts that exclusively affects the internal residues of a protein. In these cases, the alternative isoform’s reading frame is shifted due to one splicing event, but then shifts back into register of the reference frame due to a second, independent splicing event. We refer to these events as “snapback” frameshifts, as there is a return back to the original reading frame. Snapback frameshifts have been previously observed, such as in the gene *HSF4*, which produces internally frameshifted isoforms that have been demonstrated to exert different regulatory effects (Tanabe et al. 1999), but the snapback phenomenon generally speaking has not been systematically described. Within GENCODE, we found 118 examples of such snapback isoform across 95 genes, including *FAS*, and *PLEKHJ1* (**Figure 4D**, **Supplementary Table S8**). What is notable about these cases is that the same underlying genomic sequence encodes different amino acid residues, and genetic mutations could lead to two different residue changes depending on the isoform.

### Analysis of C-terminal variations

C-terminal changes make up 28% (12,241 of 44,326) of reference-alternative pairs, corresponding to 6,138 genes (**Supplementary Table S9**).

To break down the sources of C-termini variability, we found it useful to distinguish the most direct preceding cause of altered C-termini. In principle, all C-terminal changes must arise from an upstream splicing event that influences the termination codon used (notwithstanding post-translational cleavage events). However, such changes could be further classified. The altered C-terminus could arise from alternative terminal (i.e., last) exons, each harboring a different stop codon, so that the splicing event more or less directly influences the stop codon availability (i.e., direct splice-driven events). In other instances, C-terminal changes could arise from a somewhat indirect relationship to the splice event, such as when a splicing event causes a translational frameshift, in effect, “revealing” in the other frame a new stop codon that is now decoded by the ribosome (i.e., frameshift-driven events) (**Supplementary Table S9**).

In direct splice-driven events, the stop codon availability is dictated by the actively transcribed regions that contain the stop codon. We find this scenario for 72% (8,877 of 12,241) of all C-terminal variations (**Figure 5A**). These variations can be further classified based on the pattern of splicing at the C terminus. The first pattern involves an exon extension into the intron region, introducing a premature stop codon. These exon extensions introduce termination, or “EXIT”, make up 3,498 (39.4% of 8,877) cases (**Figure 5B**). The second pattern involves usage of alternative terminal coding exons or “ATE”, making up 3,301 (37.1% of 8,877) of cases (**Figure 5B**). Overall, EXIT and ATE changes lead to a shorter C-termini in the alternative isoform (distribution shown in **Figure 5C**).

**Figure 5:**
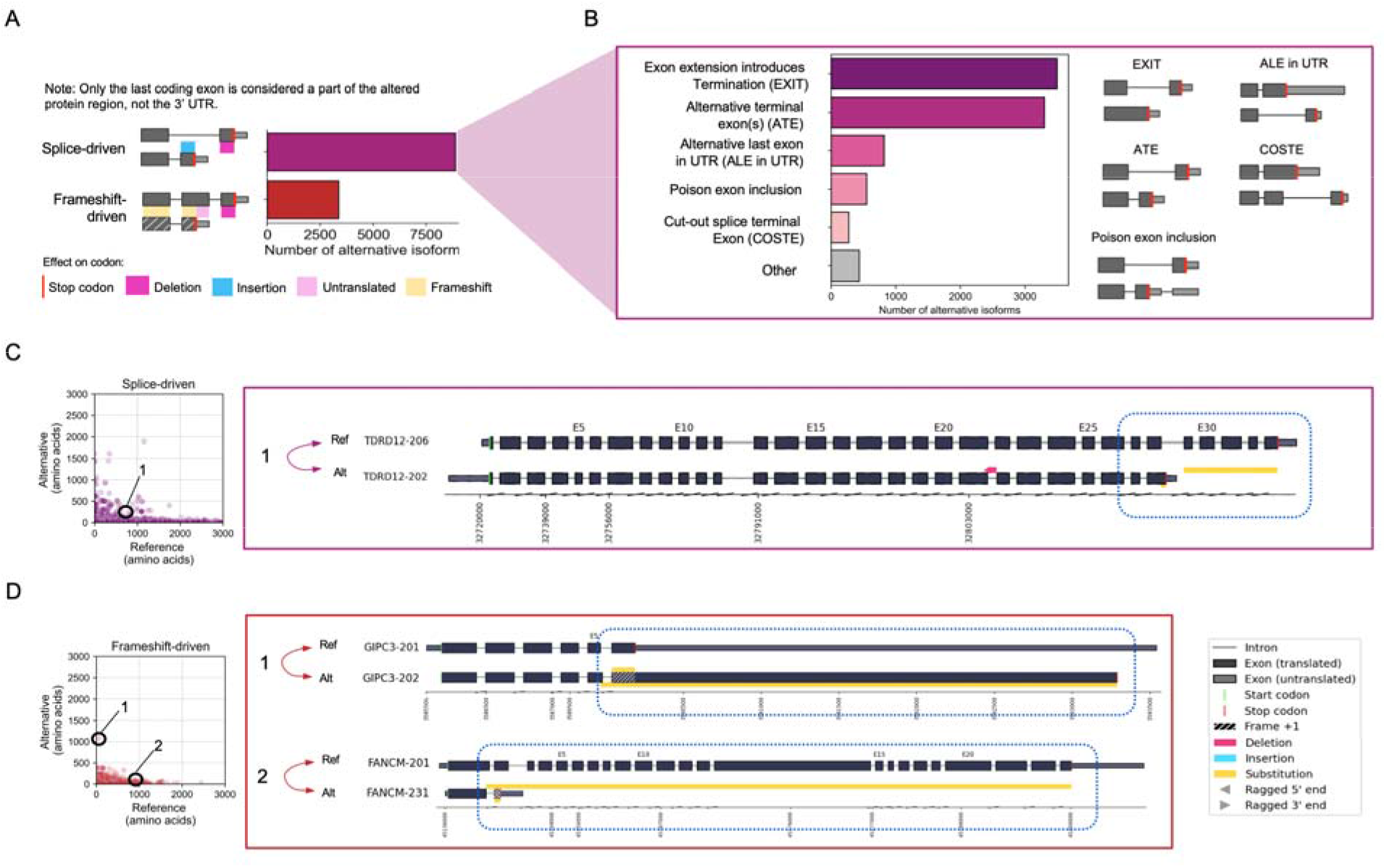
Analysis of alternative C-terminal protein sequences in GENCODE v42. (A) Frequency of alternative C-terminal categories based on splicing or frameshifts being the primary driving factor. (B) Distribution of the frequencies for various splice-driven patterns. (C) Scatterplot of the length of splice-driven C-terminal variation in the reference versus alternative. An example of this category is observed in the *TDRD12* gene. *TDRD12* undergoes splice-driven alteration causing an alternative terminal exon in TDRD12-202 to harbor the stop codon (D) Scatterplot of the length of frameshift-driven C-terminal variation in the reference versus alternative. Biosurfer plot examples of frameshift-driven category illustrated in *GIPC43* and *FANCM* genes.

EXIT versus ATE reflect how different “modes” of spliceosome regulation could lead to distinct C-terminal consequences. In EXIT, the reference-containing donor splice site fails to be spliced in the alternative isoform, leading to partial or full intron retention. An example of EXIT is shown in the right panel of **Figure 5C** for the pair, TDRD12-206 and TDRD102-202. In ATE, on the other hand, the spliceosome catalyzes splicing at one of two splice site acceptor sites, influencing terminal exon identity and thus stop codon used. A well-known pattern of ATE are poison exons, a mechanism by which inclusion of an exon leads to a premature termination codon that either elicits nonsense mediated decay or generates a truncated protein product (Carvill and Mefford 2020). Poison exons are evolutionarily conserved and likely play a role in downregulating gene expression (Lareau et al. 2007). We found 550 (6.2% of total 8,877) cases of potential poison exons. Other ATE patterns include one in which the alternative last exon of the alternative isoform resides in the UTR region of the reference isoform, suggesting that such sequences in the 3’ UTR could dually code both transcript and protein functional elements. We found 819 (9% of 8,877) cases of such alternative last exon in UTR (ALE in UTR) (**Figure 5B**). A third ATE variation, referred to as a cut-out splice terminal exon (COSTE), the 5’ end of the last exon is shared, but the alternative isoform utilizes a splice site that skips over the remaining portion of the last exon in the reference, thereby creating a different last exon not found in the original reference. We identified 273 cases of this pattern (3% of 8,877) (**Figure 5B** and **Supplementary Figure S5**).

Frameshift-driven events influence stop codon usage somewhat indirectly through shifts in the translational reading frame. In such cases, a splice-induced reading frame shift causes all downstream codons to be read in a different frame and stop codons are “revealed”, or decoded, by the ribosome. We found 3,364 (28% of 12,241) cases of frameshift-induced C-terminal changes (**Figure 5A**, Examples are shown in **Figure 5D**, pairs, GIPC3-201 and GIPC3-202, along with FANCM-201 and FANCM-231). Generally speaking, frameshifts lead to a dramatic shortening of the C-terminal region in the alternative isoform; however, we found 549 cases (16% of 3,364) in which the C-terminal region is longer in the alternative versus the reference isoform (**Figure D**).

We also observed across all frameshift-driven events a depletion of isoform pairs in which a large portion of the reference isoform (e.g., 2,000 AA or longer) is truncated due to a frameshift in the alternative isoform (**Figure D**), a trend not observed in an experimentally predicted proteome (see section below and **Supplementary Figure S9B**), likely representing gene annotation decisions, as dramatically truncating frameshift events would lead to predicted NMD and filtered out or reassigned an NMD biotype (Harrow et al. 2012).

### Characterization of altered protein regions across a long-read predicted proteome

The process of defining the reference proteome heavily draws from sources of experimental evidence such as deeply sequenced cell and tissue types, and the protein isoform sequences represent an aggregate model of the human proteome. Therefore, to characterize potential isoform related protein variability in a specific biological condition, we employed a “long read proteogenomics” pipeline (Kreitzer et al. 2013), generating a proteome predicted from long-read RNA-seq transcript sequences collected from a karyotypically normal human stem cell line (WTC-11). Using Biosurfer, we characterized the landscape of protein isoforms in the WTC-11 proteome and found 44,962 protein isoform pairs across 10,144 genes (**Supplementary Table S10**, p-block and c-block outputs in **Supplementary Table S11 and S12**). Assigning the highest expressed transcript as the “reference”, we defined 53,915 altered protein regions (**Supplementary Figure S6**). Compared to the GENCODE analysis, similar trends were observed for N-terminal (**Supplementary Figure S7**) and internal region variation (**Supplementary Figure S8**, snapback isoforms in **Supplementary Table S13**). Besides these similar trends, the experimental proteome returned a higher number of C-terminal variations that involved intron retention events (EXIT event type, see **Figure 5B**) as compared to GENCODE isoforms, matching earlier findings from EST and cDNA data (Nakao et al. 2005)(Modrek et al. 2001)(**Supplementary Figure S9**).

## DISCUSSION

To study the functional impact of alternatively spliced protein isoforms, it is critical to track precise differences in protein isoform sequences and link such variations to the upstream explanatory mechanisms. However, it is challenging to systematically characterize the full interplay between genomic and proteomic variations, which hinders discoveries of novel biological variations represented in a long-read RNA-seq dataset. We developed Biosurfer, a computational approach, available as a Python package, that systematically extracts protein isoform sequence variations while maintaining the explicit links to their underlying transcriptional and post-transcriptional mechanisms.

To demonstrate the utility of Biosurfer, we characterized protein isoform differences across an annotated (GENCODE) and long-read RNA-seq predicted proteome. Using Biosurfer’s interlinked transcript, codon, and protein data structures, we determined the upstream mechanisms explaining isoform alterations, uncovering surprising complexity. First, we confirmed past observations of alternative transcription underlying most N-terminal variations (Reyes and Huber 2018), and most internal protein sequence differences arising from single splicing events, such as exon skipping (Wang et al. 2008). However, our study goes beyond these known trends. Upon detailed tracking of codons flanking these altered protein regions, we found distinct split codon patterns that change the encoded amino acid residue identity and thus contribute to variation of AA residues. We also found an unusual frameshift pattern that involves successive reading frame shifts that leads to a change in protein regions that is entirely internal to the protein, referred to here as “snapback” frameshifts. And last, C-terminal differences are primarily splice-driven or frameshift-driven, and highly truncated alternative isoforms from frameshifts are underrepresented in GENCODE annotations but not in an experimentally proteome predicted from long-read RNA-seq data.

Biosurfer’s focus is distinct among the landscape of isoform tools, but the panoply of tools, steadily growing, may cause confusion as to the precise aspect of isoform biology being characterized by each tool. Today, many published tools process short read or long-read RNA sequencing data for the purpose of assembling full-length transcripts (Trinity)(Haas et al. 2013), discover novel transcript variations (Li et al. 2018), or quantifying splice events or entire isoforms (MISO, rMATs, RSEM, Kallisto)(Katz et al. 2010; Li and Dewey 2011; Shen et al. 2014; Bray et al. 2016). Other tools classify transcript exonic structures for novel transcripts derived from long-read RNA-seq analysis (e.g., SQANTI, FLAIR, Bambu)(Tardaguila et al. 2018; Tang et al. 2020; Chen et al. 2023). Tools like SUPPA deconstruct full-length transcriptomes into individual splicing events (Alamancos et al. 2015). Collectively, these tools analyze properties of transcripts, but with less focus on the protein effects(Altschul et al. 1990). On the other hand, tools for comparison of proteins incorporate protein sequence alignment (e.g., ClustalW, BLAST) (Chenna et al. 2003), but such alignments are disconnected from information about the underlying genome. Indeed, there are several protein-to-genome alignment algorithms (e.g., Exonerate, miniprot)(Slater and Birney 2005; Li 2022), as well as, more recently, methods to align proteins with some knowledge of the underlying exonic and genomic locations of residues, such as the Mirage tool (Nord et al. 2018; Hanimann et al. 2022; Nord and Wheeler 2023).

Related to the comparison of proteins, we had to designate a “reference” isoform, although the biological role of most isoforms are unknown (Yang et al. 2016; ReixachslSolé and Eyras 2022), and thus representative isoforms are chosen depending on the assumptions and goals of the research community (The UniProt Consortium 2017; Pozo et al. 2022).

Biosurfer analyses rely on user-defined protein isoforms. Only canonical start and stop codons are assumed, unless non-canonical sites are annotated in a reference proteome (Mudge et al. 2022) or the user. Determination of the biologically relevant ORF remains an ongoing challenge. Many ORF callers like transdecoder, CPAT, GMST and others predict ORFs, relying on heuristics and common features of translation, which may not be the rule in every case. Currently, the prediction of proteins from deep coverage long-read RNA-seq datasets rely on heuristics, such as prioritizing ORF from an alternative isoform that shares the same start AUG codon with the reference, or selection of the most 5’ proximal AUG (Tang et al. 2020; Miller et al. 2022), whereas others have developed computationally efficient scoring strategy that ranks more highly the ORFs with highest protein similarity to the reference (Varabyou et al. 2022, 2023). To provide more reliable ORF annotations, experimental approaches like Ribo-Seq demarcate novel coding regions, including sites of non-canonical translation, which might be information that could be incorporated in proteogenomic workflows (Mudge et al. 2022; Leblanc et al. 2024).

Our first version of Biosurfer proposes a new framework for detailed comparison of protein isoforms, a first step towards inferring function. Further versions of this tool could map functional elements, such as structural domains, active sites, post-translationally modified sites, or protein interactions, onto the altered protein regions, similar to the functionality of tappAS, isoTV, and DIGGER, as well as other tools (de la Fuente et al. 2020; Annaldasula et al. 2021; Louadi et al. 2021). Furthermore, given the link between genomic coordinates and effects on protein isoforms, Biosurfer could capture the impact of coding or splice-modifying genetic variants as carried through the lens of complex transcript and protein variations, which might increase the accuracy of predicted genetic effects in ancestry- or patient-specific populations (Rivas et al. 2015)(Cummings et al. 2017)(Yamaguchi et al. 2022)(Glinos et al. 2022). With increasing appreciation for population scale diversity and the pan-genome, which, carried forward, gives rise to a corresponding “pan-proteome”, bioinformatic pipelines should be designed to produce automated results of all possible variations arising from newly sequenced sample. Ultimately genotype could be associated to the full repertoire of proteoform diversity (Aebersold et al. 2018), especially as proteomics approaches continues to capturing a greater swath of the protein isoform space (Sinitcyn et al. 2023).

## DATA ACCESS

All raw data has been previously published and is described in the **Methods**.

## ACKNOWLEDGMENTS

This work was supported by the National Library of Medicine (R01-LM014017) to G.M.S. and D.K.

